# Phoronids and their tubes harbor distinct microbiomes compared to surrounding sediment

**DOI:** 10.1101/2024.05.28.596327

**Authors:** Cassandra L. Ettinger, Jonathan A. Eisen

## Abstract

Phoronids are a phylum of animals with only ∼12 described species, all of which are marine filter feeders that build external tubes for shelter and produce chemical deterrents against predators. Many tube-building invertebrates host distinct microbial communities and even have obligate symbionts for survival in sulfur-rich marine sediments. However, the microbiome of phoronids has only recently begun to be described. To address this, we surveyed the composition of the microbiome of the phoronid, *Phoronopsis harmeri*, using 16S rRNA gene amplicon and metagenomic sequencing. We found that the phoronid microbiome was dominated by members of the orders Campylobacterales, Desulfobulbales, and Desulfobacterales. We also found that the microbiomes of tubes and phoronids were less diverse than that of surrounding sediment, and that the microbiomes of phoronids, tubes and surrounding sediment were all distinctly structured. Based on analysis of metagenomic data, and even though we were only able to recover low quality MAGs of abundant taxa, we found preliminary evidence that taxa associated with phoronids and their tubes likely participate in sulfur cycling pathways. Future work should perform more robust metagenomic sequencing and chemical analysis to assess if there is a link between known phoronid chemical defenses and microorganisms. Overall, this study provides foundational insight into the microbial communities associated with phoronids and these initial findings suggest that these communities may play an important role in sulfur cycling in marine sediments.

## Introduction

*Phoronopsis harmeri* [1], previously known as *P. viridis* [2], is a phoronid found in several small bays in central California, including Bodega Harbor. Phoronids are lophophorate organisms, aquatic invertebrates that have a specialized feeding structure made of tentacles, that live in marine sediment and build external tubes for shelter. Thus, phoronids are like other marine worm-like organisms that serve as ecosystem engineers through creation of dwellings such as tubes, mounds or other structures that alter sediment stability, enhance solute exchange and thus create distinct local environments [3–16]. *P. harmeri* is a digging phoronid and has an organic tube consisting of particles of the surrounding sediment [17]. These tubes consist of an inner organic cylinder made of thin films whose fibers form a thick net and an external layer [18]. Phoronid tubes can be 1 to 3 mm in diameter and up to 20 cm long, about twice the length of the phoronid [18,19]. While typically found inside their tubes, phoronids are not physically attached to their tubes, and in order to filter feed, they extend their green colored lophophores above the sediment [14,20,21]. Phoronid tubes can occur at very high densities and can lead to increased diversity of infaunal community members [19].

Studies of other marine ecosystem engineers, including polychaetes (e.g., *Escarpia* sp. [22], *Diopatra* sp. [23], *Neanthes* sp. [24]), tube worms (e.g., *Riftia* sp. [25], *Lamellibrachia* sp. [26]) and shrimp (e.g., *Upogebia* sp. [27]), have described important roles for the microbiome in host-environment interactions including metal resistance and sulfur cycling [28–30]. Often the microenvironment of the tube or burrow created by these marine organisms is found to have a distinct microbiome from surrounding sediments [22,23,28,31]. For example, blue mud shrimp burrows [27] introduce oxygen into local sediment which supports biogeochemical zonation around the burrow and may promote growth of sulfur cycling microbes. Free-living sulfur oxidizers and reducers have been reported to be enriched in association with marine ecosystem engineers and their respective structures [27,31–34]. Bivalves in particular form tight symbiotic associations with sulfur oxidizing bacteria which provide the bivalve with primary production products, like sugars, in exchange for hydrogen sulfide or oxygen [35,36]. It has been suggested that the tube of phoronids may serve as a physical barrier against hydrogen sulfide and allow for a similar zone of oxygenation [37], and while bacteria have recently been observed to line the inside of phoronid tubes with microscopy [18], to date, no symbiotic or free-living sulfur cycling microbes have been confirmed to associate with phoronids.

In addition to tube-building, the phoronid, *P. harmeri,* is known to produce a chemical deterrent that is unpalatable to predators [20]. While all parts of the phoronid exhibit some predator deterrence, this feature is most concentrated in its green lophophore [20]. In *P. harmeri,* the chemical compound responsible has not yet been successfully isolated, likely due to high volatility or reactivity with air [20]. While many other structure-forming infaunal organisms also produce secondary metabolites to deter predators, historically few efforts have successfully isolated the responsible compounds [38–45]. Recent studies of other marine organisms (e.g., sea slugs, sponges) have attributed the origin of their chemical defense to symbiotic microbes [46,47]. Microscopic anatomy studies of phoronids have visualized bacteria inhabiting the lophophore organs of adults [48,49], leading us to speculate on whether there is a role for microbes in the production of the chemical defenses of phoronids and other structure-forming infaunal organisms. While a recent study described the bacterial microbiome of a different phoronid species in British Columbia and suggested that the tube was the primary driver of overall microbiome composition [50], the microbiome of phoronids and their tubes relative to surrounding sediment has yet to be comprehensively characterized.

In this foundational study, we use sequencing and analysis of 16S rRNA gene amplicons and metagenomes to achieve the following goals (i) survey the overall microbiome composition of the phoronid, *P. harmeri,* (ii) assess whether the microbiome differs between phoronids, tubes and surrounding sediment, and (iii) generate metagenome-assembled genomes (MAGs) in order to hypothesize functional roles for the dominant microbial associates of phoronids.

## Methods

### Sample Collection

Marine invertebrates were sampled as accidental bycatch during a previous study of the mycobiome of the seagrass, *Zostera marina [51]*. In that study, individual *Z. marina* and adjacent sediment, as well as unvegetated sediment, were cored using a modified 2.375 inch diameter PVC pipe. In total, three vegetated and one unvegetated core contained phoronids as bycatch. All four of these cores were collected from one site in Bodega Bay, CA, Westside Point (GPS: 38°19′10.67″N, 123° 3′13.71″W) in August 2016. A sediment sample was collected from each core using an 11 mm diameter straw. Cores were transported on dry ice to the lab and bycatch was removed using flame-sterilized sterile tweezers. Individual phoronids were extracted from their associated tube and both phoronid and tube were placed into separate sterile 1.5 mL tubes. In total, four sediment samples, 13 phoronids (*Phoronopsis harmeri*), and 12 tubes were opportunistically collected. This included 11 paired sets of phoronids and their respective tubes. All samples were stored at -80 °C until processing.

### Molecular methods

Samples were randomized prior to extraction using a random number generator. All marine invertebrate samples were surface cleaned and sterilized as in Rattray et al [52]. Briefly, this involved rinsing samples in autoclaved water, then immersing samples in 95% ethanol for 5 minutes, followed by a final rinse in autoclaved water. DNA was extracted from surface sterilized samples with the ZymoBIOMICS DNA Mini kit (Zymo Research, Irvine, CA) following the manufacturer’s protocol with samples bead beaten on the homogenize setting for 1 min using a BioSpec Products mini-bead beater. DNA was also extracted from three no-sample added (negative), three autoclaved water (negative), three 95% ethanol (negative), three 70% ethanol (negative) and three ZymoBIOMICS Microbial Community Standard (positive) controls (Zymo Research, Irvine, CA).

### Sequence generation

For high-throughput amplicon library preparation and sequencing, samples were randomized and placed in 96-well plates to be sent to Zymo Research, Inc. via their ZymoBIOMICS Service. The 16S ribosomal RNA (rRNA) gene was amplified using the Earth Microbiome Project (EMP) 515F (Parada) and 806R (Apprill) primer set [53,54]. Libraries were produced using the Quick-16S™ NGS Library Preparation Kit (Zymo Research, Irvine, CA). PCR products were quantified with qPCR and pooled together based on equal molarity. The pooled library was cleaned with Select-a-Size DNA Clean & Concentrator^TM^ (Zymo Research, Irvine, CA, United States), then quantified with TapeStation (Agilent, Santa Clara, CA, United States) and Qubit (Invitrogen, Carlsbad, CA, United States). The final library was sequenced on an Illumina MiSeq (Illumina, Inc., San Diego, CA, United States) with a v3 reagent kit (600 cycles) to generate 300 bp paired-end reads. The raw sequence reads generated for this 16S rRNA gene amplicon project were deposited at GenBank under BioProject ID PRJNA1016428.

Two paired samples (one phoronid [511], and one tube [512]) were chosen for metagenomic sequencing. DNA quality was checked on a 1.5% E-Gel (Thermo Fisher Scientific, Waltham, MA, United States). DNA was provided to the UC Davis Genome Center DNA Technologies Core for library preparation and sequencing as part of a shared multi-project sequencing run of nine samples. DNA libraries were sequenced on an Illumina MiSeq to generate 250 bp paired-end reads. The raw metagenomic sequence reads generated for this project were deposited at GenBank under BioProject ID PRJNA1016428.

### 16S rRNA gene amplicon processing

Primers were removed from reads using cutadapt v. 2.3 [55]. Reads were processed with DADA2 in R with the following parameters maxEE=c(2,2), truncLen=c(190, 190), and truncQ=2 [56–58]. Using DADA2, Reads were then denoised and merged to generate amplicon sequence variant (ASV) count tables. Chimeric sequences were identified and removed using removeBimeraDenovo by “consensus” (1.22% of ASVs).

After removing chimeras, samples had a mean read count of 50,507 (range: 27,899 - 74,130). Taxonomy was inferred for ASVs using the RDP Naive Bayesian Classifier algorithm with the SILVA (v. 138) [59–61]. ASVs were aligned using DECIPHER [62,63]. A neighbor-joining tree was then constructed as a starting tree, and fit to a GTR+G+I (Generalized time-reversible with Gamma rate variation) maximum likelihood phylogeny using phangorn [64]. The resulting phylogeny was then rooted using a eukaryotic outgroup with APE [65].

To identify possible contaminants, we used decontam’s prevalence method with a threshold of 0.5, which will identify ASVs with a higher prevalence in negative controls than in true samples [66]. This threshold identified 113 possible contaminant ASVs which were removed from the final dataset. Negative and positive controls were subsequently removed at this point in the analysis. Further, all ASVs with taxonomic assignments in the Eukaryota or an unclassified kingdom, as well as to chloroplasts or mitochondria were removed. The resulting count table contained 4,295 ASVs.

Count tables were normalized for sequencing depth by subsetting without replacement to 10,000 sequences per sample. Rarefaction depths were chosen to balance rarefaction curve saturation with the minimum library size needed to maintain all samples.

### 16S rRNA gene amplicon data analysis

Relative abundance, alpha and beta diversity analyses were assessed using the tidyverse [67], ggplot2 [68], vegan [69] and phyloseq [70] packages in R. We visualized the relative abundance of the microbiome across sample types (phoronids, tubes, sediment) by transforming rarified read counts to proportions and collapsing ASVs into taxonomic families. To better enable interpretation, we only visualized families or orders that represented a summed mean proportion of greater than 5% relative abundance across sample types.

To assess differences in alpha diversity across sample types, we calculated the Shannon index and then tested for significant differences using Kruskal-Wallis (KW) tests with 9,999 permutations. When the KW test resulted in a rejected null hypothesis (*P*□*<*□0.05), we further applied *post hoc* Dunn tests and corrected the resulting *p*-values using the Benjamini-Hochberg method.

To assess differences in beta diversity across sample types, we subsampled to the desired rarefaction depth using rarefy_even_depth from phyloseq across 100 random iterations, calculated the Weighted Unifrac dissimilarity for each iterative dataset using the distance function from phyloseq, and then averaged across all iterations to generate an average dissimilarity matrix for the dataset. We visualized the average dissimilarity matrix as a principal coordinate analysis (PCoA) plot using the ordinate function in phyloseq. We tested for significant differences in beta diversity centroids (i.e., means of each group) across sample types using permutational multivariate analyses of variance (PERMANOVAs) with 9,999 permutations and by = “margin” using the adonis2 function. Vegetative status (i.e., with or without seagrass) was included as a covariate in the PERMANOVA model. *Post hoc* pairwise PERMANOVAs were performed to assess pairwise differences using the pairwise.adonis function from the pairwiseAdonis package [71] with *p*-values corrected using the Benjamini-Hochberg method. We also tested for significant differences in dispersion (i.e., variability) using the betadisper and permutest functions from the vegan package with 9,999 permutations.

### Metagenomic read-based analysis

Adaptor removal and quality filtering of metagenomic reads was performed using BBDuk v. 38.90 [72] with the following parameters: ktrim=r k=23 mink=11 hdist=1 tpe tbo qtrim=rl trimq=10 maq=10. Taxonomic classification of quality-controlled reads was performed using Kraken2 v. 2.1.2 [73] against the NCBI non-redundant nucleotide database (downloaded May 3, 2023). Bracken v. 2.5 [74] was then run on the Kraken2 outputs to estimate the number of reads associated with each species in each individual sample. ETE3 [75] was used to obtain the full taxonomic lineage string of each identified species from NCBI.

Resulting Bracken count tables were imported into R for downstream analysis using the tidyverse [67], ggplot2 [68], vegan [69] and phyloseq [70] packages in R. Bracken counts were transformed into proportions. We visualized the relative abundance of the microbiome across metagenomic samples (one phoronid, one tube). To better enable interpretation, we only visualized families or orders that represented a summed mean proportion of greater than 5% relative abundance across samples.

### Metagenome-assembled genome (MAG) analysis

Quality filtered reads were coassembled using MEGAHIT v1.2.9 [76]. The anvi’o v.7.1 pipeline was then used to identify and assess metagenome-assembled genomes (MAGs) [77]. Briefly, we first calculated read coverage against the coassembly using bowtie2 v. 2.4.2 and samtools v. 1.11 [78,79]. Then, we constructed a contig database for the coassembly using ‘anvi-gen-contigs-database’ and predicted genes using Prodigal v. 2.6.3 [80] and single-copy genes for bacteria [81] using HMMER v. 3.2.1 [82]. We then assigned taxonomy to genes using Kaiju v. 1.8.2 with the NCBI BLAST non-redundant protein database nr including fungi and microbial eukaryotes v. 2020-05-25 [83].

Anvi’o profiles were constructed for each individual sample for contigs > 1 Kb using ‘anvi-profile’ with the ‘--cluster-contigs’ option. Individual sample profiles were subsequently merged using ‘anvi-merge’. Contigs were automatically binned into MAGs using ‘anvi-cluster-contigs’ with four different algorithms, CONCOCT v. 1.1.0, BinSanity v.0.5.4, MaxBin v. 2.2.7 and METABAT v. 2.2.7 [84–87]. An optimized, non-redundant set of MAGs was then calculated using DASTOOL v. 1.1.2 [88]. Completeness and contamination of MAGs were assessed using ‘anvi-summarize’ and CheckM2 [89]. Inspection of MAGs was performed using ‘anvi-refine’ and manual refinement was performed as needed. Taxonomy was assigned to MAGs using GTDB-Tk v.2.2.6 [90], which uses both phylogenetic placement and average nucleotide identity (ANI) to make taxonomic assignments. METABOLIC v.4.0 was run using ‘METABOLIC-G.pl’ on both the coassembly and individual MAGs to identify genes related to sulfur cycling processes [91]. Only MAGs with ≥ 25% completion based on CheckM2 estimates are reported here.

## Results

### Phoronid microbiome is dominated by members of the order Campylobacterales

Analysis of the 16S rRNA gene amplicon data revealed that the taxonomic orders with the highest mean relative abundance associated with phoronids were the Campylobacterales, Verrucomicrobiales, Desulfobulbales, and Desulfobacterales (Figure 1A, Figure S1). For tubes, the orders with the highest average relative abundance were Campylobacterales, Desulfobulbales, Desulfobacterales, and Flavobacteriales. Finally, the most abundant orders associated with sediment were Campylobacterales, Desulfobulbales, Flavobacteriales, and Desulfobacterales. Overall, the most relatively abundant taxonomic orders associated with all three sample types were almost identical, possibly indicating that much of the phonorid and tube associated communities may be sourced from the surrounding sediment.

**Figure 1.**
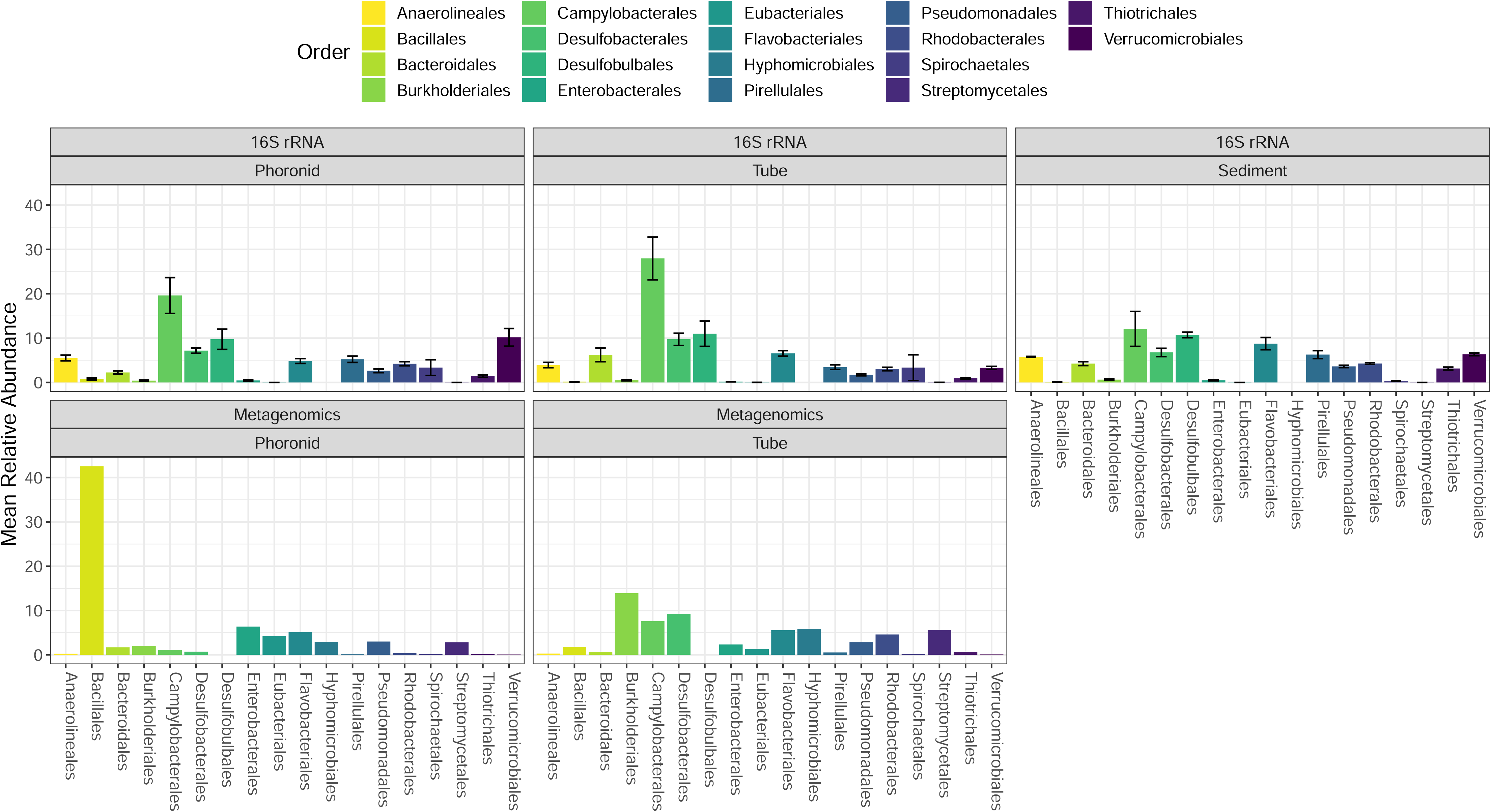
Mean relative abundance of orders associated with phoronids, tubes and surrounding sediment based on analysis of 16S rRNA gene amplicon and metagenomic data. Bar charts display the mean relative abundance of taxonomic orders for each sample type (phoronid, tube, sediment) colored by predicted order. Mean relative abundance based on 16S rRNA gene amplicon data is shown on top with bars representing standard error, with mean relative abundance of read-based Kraken2/Bracken2 analysis of metagenomic data shown on the bottom. Orders representing less than 5% mean relative abundance across datasets are not shown. Number of samples summarized as follows - Phoronid: *n_16S_* = 13, *n_Metagenomics_*= 1; Tube: *n_16S_* = 12, *n_Metagenomics_* = 1; Sediment: *n_16S_* = 4.

In comparison, a read-based analysis of the metagenomic data using Kraken2/Bracken found the taxonomic orders with the highest relative abundance associated with phoronids to be the Bacillales, Enterobacterales, Flavobacteriales, and Eubacteriales, very different from the 16S rRNA gene amplicon results (Figure 1B, Figure S2). The taxonomic orders with the highest relative abundance associated with tubes in the metagenomic read analysis (Burkholderiales, Desulfobacterales, Campylobacterales, and Hyphomicrobiales) were largely similar to the 16S rRNA gene amplicon results. There are several possible reasons for discrepancies between the metagenomic and the 16S rRNA gene amplicon data including primer bias [92,93], different naming conventions between databases [94,95], and the limitations of read-based taxonomic classifiers [96]. However, one major contributor here was likely the high-prevalence of host DNA in the phoronid metagenome, which may have limited our ability to accurately determine relative abundance of bacterial taxa. Metagenomic read analysis predicted that only 6.40% of reads were bacterial in the phoronid metagenome, while the remaining 93.09% were predicted to be eukaryotic. In comparison, analysis of the tube metagenome yielded a much higher proportion of bacterial reads (43.09%) compared to eukaryotic reads (56.11%).

### Phoronids and tubes are distinct from surrounding sediment

We compared alpha diversity and community structure between phoronids, tubes and surrounding sediment using 16S rRNA gene amplicon data analysis. We found that phoronids, tubes and surrounding sediment had significantly different Shannon alpha diversity (Figure 2A; K-W test, p < 0.05). *Posthoc* tests revealed that phoronids and tubes had similar Shannon alpha diversity (Dunn test, p > 0.05) and both were less diverse than sediment (p < 0.05). Shannon alpha diversity was not significantly different across samples from different cores, vegetation status (whether a core was from within or outside of a seagrass bed), or between phoronid-tube pairs (K-W test, p > 0.05).

**Figure 2.**
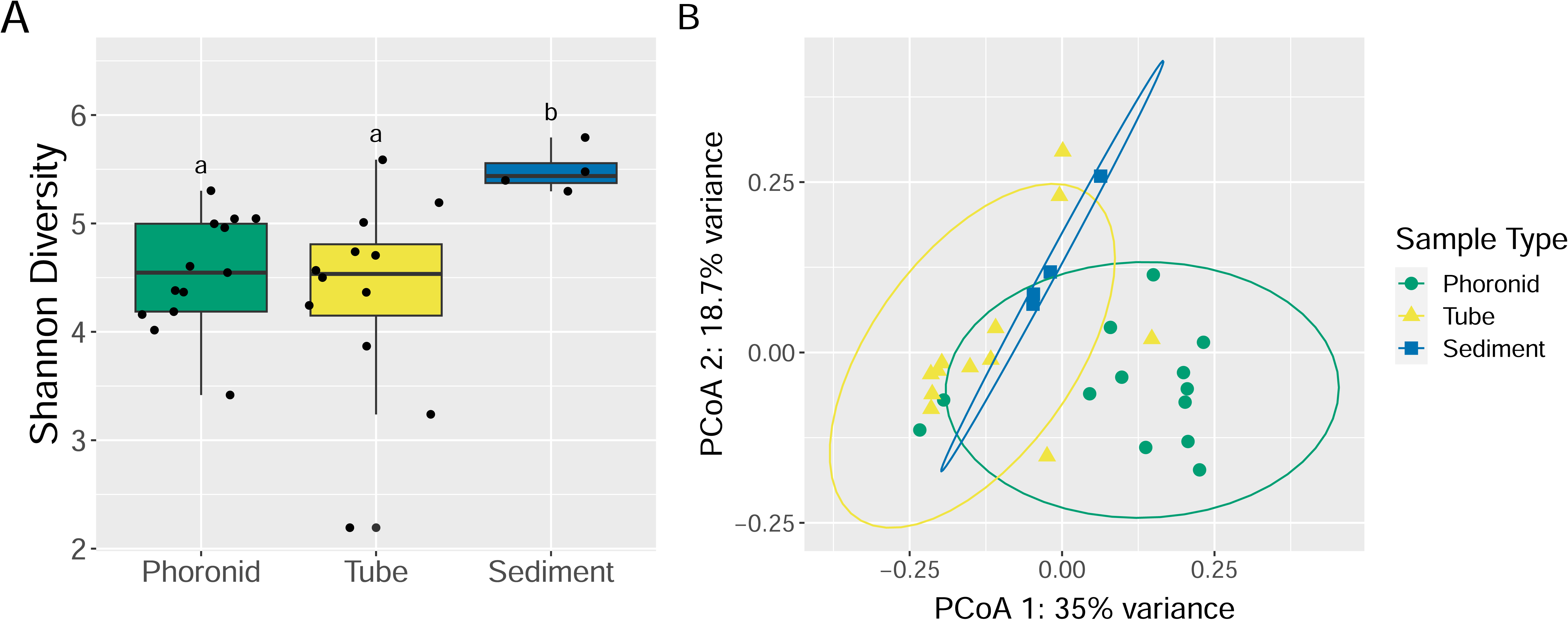
Phoronids and tubes have distinct microbial communities compared to surrounding sediment. (A) Shannon diversity was used to assess alpha diversity across sample types (phoronids, tubes, sediment). Comparisons that are significantly different from each other are notated by different letters (e.g., a vs. b). (B) Principal-coordinate analysis (PCoA) visualization of Weighted Unifrac distances shows individual samples designated as points with colors and shapes representing sample type: phoronids (green circle), tubes (yellow triangles), and sediment (blue squares). Ellipses represent the 95% confidence interval around the centroid of each group. Number of samples summarized in (A) and (B) as follows - Phoronid: *n* = 13; Tube: *n* = 12; Sediment: *n* = 4.

Phoronids, tubes and surrounding sediment also had significantly different community structure (Weighted UniFrac beta diversity) (Figure 2B; PERMANOVA; p < 0.01). *Posthoc* pairwise contrasts were all significantly different, indicating that phoronids, tubes and surrounding sediment all harbor distinctly structured communities (p < 0.05). Vegetation status also was associated with significantly different community structure (Figure S3; p < 0.01). Additionally, we found no significant differences in community variance, or dispersion, in terms of sample type or vegetation status (betadisper; p > 0.05).

### Recovery of low-quality metagenome-assembled genomes and sulfur cycling genes

Despite limitations due to prevalence of host DNA in the phoronid sample (∼93% represented eukaryotic reads), low sample size and sequencing depth, which precluded us from obtaining high (≥ 90% complete) or medium quality MAGs (≥ 50% complete) [97], we were still able to generate and report here on five low quality draft MAGs associated with phoronid tubes that were ≥ 25% complete (CheckM2; Table 1, Table S1). The taxonomic identities of the five low quality MAGs largely reflect the dominant taxonomic orders and families of phoronids and tubes found in the 16S rRNA gene amplicon data (i.e., Campylobacterales, Desulfobacterales, and Flavobacteriales). As a result of evolutionary novelty, incompleteness, or both, none of these MAGs was placed by GTDB-Tk into a species level taxonomic assignment with several of the MAGs only able to be placed at higher taxonomic levels (i.e., class, family).

**Table 1.**
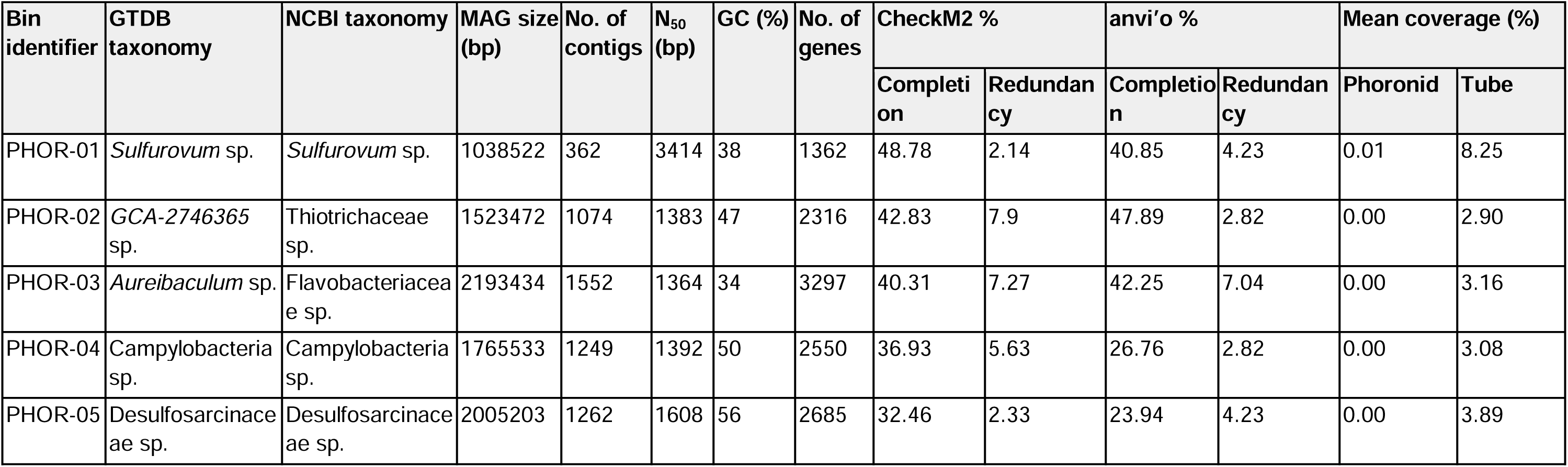
Summary of genomic features of five draft MAGs. Genomic features are summarized for each MAG, including putative taxonomic identity (GTDB and NCBI), genome size (bp), number of contigs, N50, percent GC content, number of genes, completion and contamination estimates as generated by CheckM2 and anvi’o, and mean read coverage against each metagenome. MAGs are sorted by CheckM2 completion. See Table S1 for full GTDB taxonomy.

Given the importance of sulfur cycling in marine ecosystems, we used METABOLIC to survey the sulfur cycling capacity of the metagenome co-assembly overall and the five low quality MAGs individually. Results from running METABOLIC on the co-assembly suggest that a majority of genes needed for the sulfur cycle are present in the microbial community associated with phoronids and tubes (Figure 3, Figure S4). Further, while we only have an incomplete picture of the genes present in the MAGs, METABOLIC found that some of the MAGs, in particular PHOR-02, PHOR-04, PHOR-05, had key genes involved in a subset of these pathways. With PHOR-02 likely capable of sulfite oxidation, sulfate reduction and sulfite reduction, PHOR-04 capable of sulfur oxidation, and PHOR-05 capable of sulfite reduction. Genes involved in other key sulfur cycling pathways like sulfide oxidation, thiosulfate oxidation and thiosulfate disproportionation while not detected in the MAGs, were detected in the metagenomic co-assembly.

**Figure 3.**
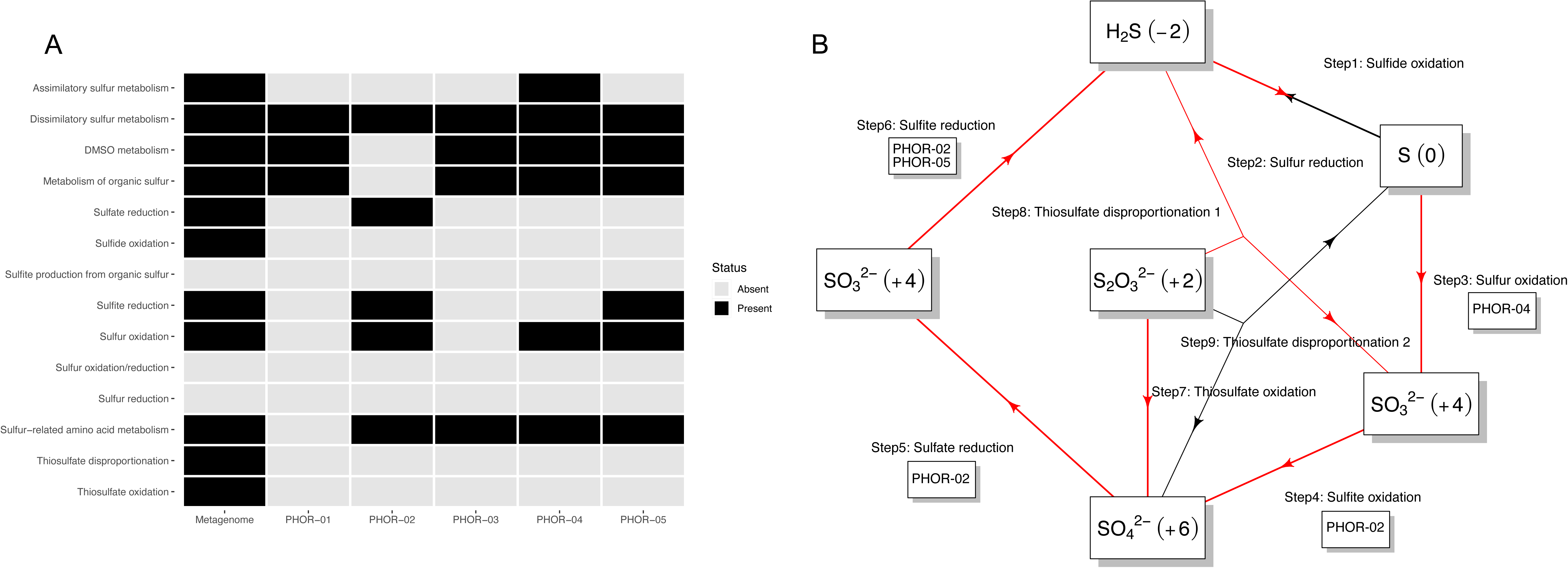
Predicted sulfur metabolism profiles. (A) Heatmap showing whether the metagenomic coassembly or individual MAGs had any genes detected by METABOLIC, which predicts metabolic and biogeochemical functional traits of input genomic data, related to each of the different sulfur cycling metabolisms and processes. A gene-level heatmap can be found in Figure S4. (B) Summary scheme of the sulfur cycle at the community scale. Each arrow represents a single step within the cycle. Red arrows indicate that the METABOLIC found the required genes for a step in the metagenomic coassembly, while black arrows indicate missing steps. Under each step is a box displaying the identifier(s) of any MAG(s) found by METABOLIC to have the genes needed for that step.

## Discussion

### Phoronid microbiome is similar to that of other marine organisms

The orders Campylobacterales, Desulfobulbales, and Desulfobacterales had the highest relative abundance in the 16S rRNA amplicon analysis of the microbiomes of phoronids, tubes, and their surrounding sediment. These orders have been reported as highly abundant members of the microbiomes of other marine organisms and their dwellings including shrimp [27,98], polychaetes [31,99,100], and tube worms [101]. Consistent with our findings, a study of a different phoronid species from British Columbia recently reported that phoronid-associated bacterial communities were distinct from the surrounding seawater, with Campylobacterota and Desulfobacterota among the organisms detected [50]. Metagenomic read-based analysis similarly identified Campylobacterales and Desulfobacterales as relatively highly abundant in the microbiome of tubes. However, the metagenomic analysis of phoronids was less consistent with the 16S rRNA amplicon analysis likely due to several factors including 16S rRNA gene primer bias [92,93], different naming conventions between 16S rRNA gene and genomic databases [94,95], database age [102,103], and methodological limitations of short read-based taxonomic classifiers [96], which can be complicated by low coverage or host-dominated metagenomic sequencing.

### Phoronid microbiome is dominated by known sulfur-cycling microbes

Sulfur cycling is a critical ecosystem process in the marine ecosystem and many marine organisms have known associations with microbes involved in the sulfur cycle [27,31–33,104]. It is thought that remineralization of a third of organic matter on the seafloor globally is facilitated by sulfate-reducing microbes [105]. The orders Campylobacterales, Desulfobulbales, and Desulfobacterales all include taxa known to be involved in sulfur cycling [106,107]. Members of the Campylobacterales, such as *Sulfurovum* sp., include known sulfur-oxidizing chemolithoautotrophs and are often reported as chemosymbionts of marine animals such as tubeworms, shrimp, snails, polychaetes, and bivalves [30,108–115]. Members of the Desulfobulbales include filamentous cable bacteria which couple sulfur oxidation and oxygen or nitrate reduction via long distance electron transfer [116,117]. Finally, members of the Desulfobacterales include sulfate reducers and oxidizers, and symbionts of gutless marine oligochaetes [118,119]. Understanding the intricate relationships between phoronids and these sulfur-cycling microbes may shed light on the potential contributions of phoronids to sediment biogeochemical cycling and provide valuable insights into understanding broader marine ecosystem dynamics.

### Tube habitat may help select for sulfur-cycling microbes from sediment

Phoronid tubes hosted a community that was distinct from both host phoronid and surrounding sediment, despite having a similar taxonomic composition to both. This pattern has also been observed in phoronids from British Columbia, where the microbiome of the tube was found to be similar to that of the whole animal, with Holt et al. suggesting the tube as the dominant niche structuring the overall phoronid-associated community [50]. Distinct tube or burrow microbiomes have been reported for shrimp [27] and polychaetes [100,120], and it is thought that these dwellings may provide conducive conditions for sulfur reduction, potentially explaining the prevalence of orders like Desulfobulbales and Desulfobacterales for which many of the known members are sulfate-reducers [7,31,120,121]. Desulfobulbales are thought to be able to tap into polychaete tubes which serve as oxygenated conduits facilitating sulfur cycling [122]. Similar findings in mud shrimp burrows suggest a role for these dwellings in providing a zone of oxygenation for sulfate-reducing microbes [27]. These results combined with observations of bacteria lining the inside of phoronid tubes [18], suggests that these tubes may be creating a microenvironment conducive to sulfur-cycling microbes, echoing the observations of other dwelling-building species.

The phoronids and their tubes also hosted less diverse microbiomes than that of surrounding sediment, which could indicate that there is some selection occurring. Host-associated drops in diversity have been reported in other studies of marine invertebrates in comparison to surrounding habitat [27,123]. However, a comprehensive analysis of marine invertebrate microbiomes found that while microbiomes were distinct from surrounding sediment, providing some evidence of host selection, there was no evidence of phylosymbiosis [123]. This suggests that local habitat is a defining driver of microbial community composition in marine species. Here we found a significant effect of vegetation status, which further supports the idea that local habit is important as seagrasses have previously reported to select for a distinct soil community compared to surrounding unvegetated sediment [124]. Similarly, Holt et al. found that the microbiome of phoronids from the same colony shifted substantially over time, further supporting the idea that local environmental conditions, rather than host identity alone, are a primary driver of community composition in phoronids [50]. These findings underscore the intricate interplay between marine invertebrates and their microbial communities in shaping local ecosystems, as well as the role of local ecosystems in shaping these host-microbe interactions.

### Functional analysis of MAGs support role in sulfur-cycling

We were able to generate five low quality draft MAGs including members of the dominant orders Campylobacteria and Desulfobacterales. These partial MAGs point towards involvement of abundant taxa and other microbial community members in sulfur cycling. We found that PHOR-02 (GCA-2746365 sp./Thiotrichaceae sp.) was likely capable of sulfite oxidation, sulfate reduction and sulfite reduction, PHOR-04 (Campylobacteria sp.) capable of sulfur oxidation, and PHOR-05 (Desulfosarcinaceae sp.) capable of sulfite reduction. However, we are likely undersampling the true functional potential of these MAGs due to their relatively incomplete status. For example, PHOR-01 (*Sulfurovum* sp.) is missing many of the genes required for different sulfur cycling processes, despite belonging to a genus that includes reported sulfate oxidizers [101,125,126]. Regardless, these partial MAGs provide initial insight into the possible function of microbes associated with phoronids and their tubes, depicting a microbial community likely involved in marine sulfur cycling. Given that microscopic evidence suggests that there are symbiotic microbes associated with phoronids [48,49], future work should improve on these MAGs to enable investigation into other functional roles of these microbes in phoronid fitness, for example, whether microbes may play a role in host chemical defense or host nutrition.

## Conclusion

While limited in scope, this work provides a foundation for understanding phoronid microbial communities. We found that phoronids and tubes were distinct in microbiome richness and community structure compared to nearby sediment. Further, we identified that phoronids and their tubes harbor distinct microbial communities. This suggests that phoronids and tubes are providing different niches for microbial community members. In summary, this work describes the microbiome of another marine tube-building organism that may have formed associations with microorganisms involved in sulfur metabolism to survive in marine ecosystems. These results also open the door to future exploration of whether there is a microbial contribution to phoronid biochemical defense against predators or host nutrition.

## Author Contributions

CLE conceived and designed the experiments, performed sampling, analyzed the data, prepared figures and/or tables, wrote and reviewed drafts of the paper. JAE reviewed drafts of the paper.

## Data availability

This 16S amplicon sequencing project and raw metagenomic sequencing reads have been deposited at GenBank under the accession no. <u>PRJNA1016428</u>. Individual MAG assemblies are available on Zenodo (https://zenodo.org/doi/10.5281/zenodo.11224923). All data and code can be found in GitHub (https://github.com/casett/PhoronidMicrobiomes) and is archived on Zenodo (https://zenodo.org/doi/10.5281/zenodo.11225270).

## Acknowledgements

We would like to thank Katherine Dynarski (ORCID: <u>0000-0001-5101-9666</u>) and Sonia Ghose (ORCID: <u>0000-0001-5667-6876</u>) for help with sample collection. We would like to thank John J. Stachowicz for use of his scientific sampling permit, California Department of Fish and Wildlife Scientific Collecting Permit # SC 4874.

