## Supplemental Material for "Phoronids and their tubes harbor distinct microbiomes compared to surrounding sediment"

**Supplemental Figures and Tables:**

**Figure S1.** Mean relative abundance of families associated with phoronids, tubes and surrounding sediment using 16S rRNA gene sequence data. Bar charts display the mean relative abundance of taxonomic families for each sample type (phoronid, tube, sediment) colored by predicted family with bars representing standard error. Families representing less than 5% mean relative abundance are not shown. Number of samples summarized as follows - Phoronid: *n* = 13; Tube: *n* = 12; Sediment: *n* = 4.

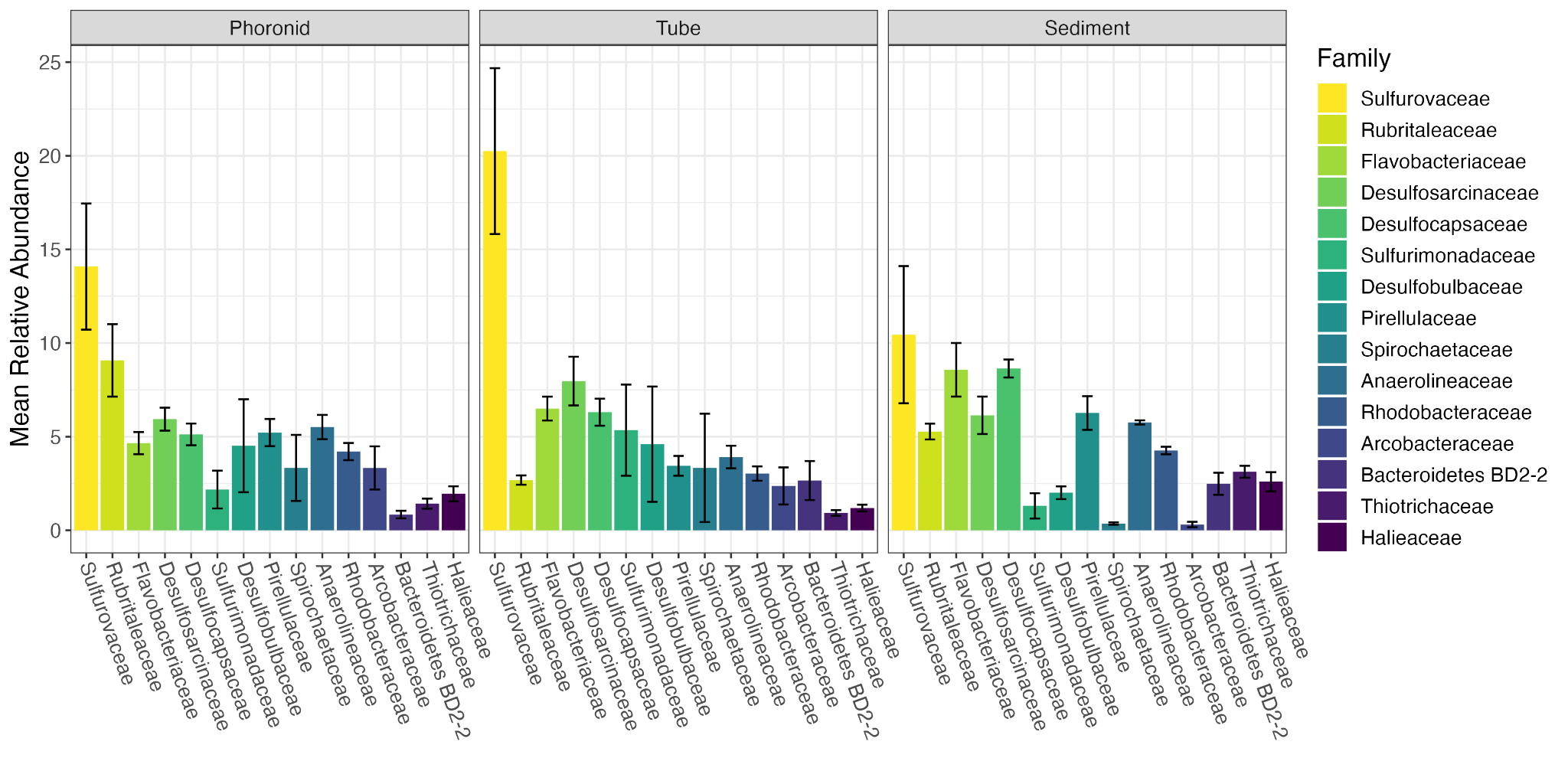

**Figure S2.** Mean relative abundance of families associated with phoronids, tubes and surrounding sediment using metagenomic data. Bar charts display the mean relative abundance of taxonomic families for each sample type (phoronid, tube) colored by predicted family. Families representing less than 5% mean relative abundance are not shown. Number of samples summarized as follows - Phoronid: *n* = 1; Tube: *n* = 1.

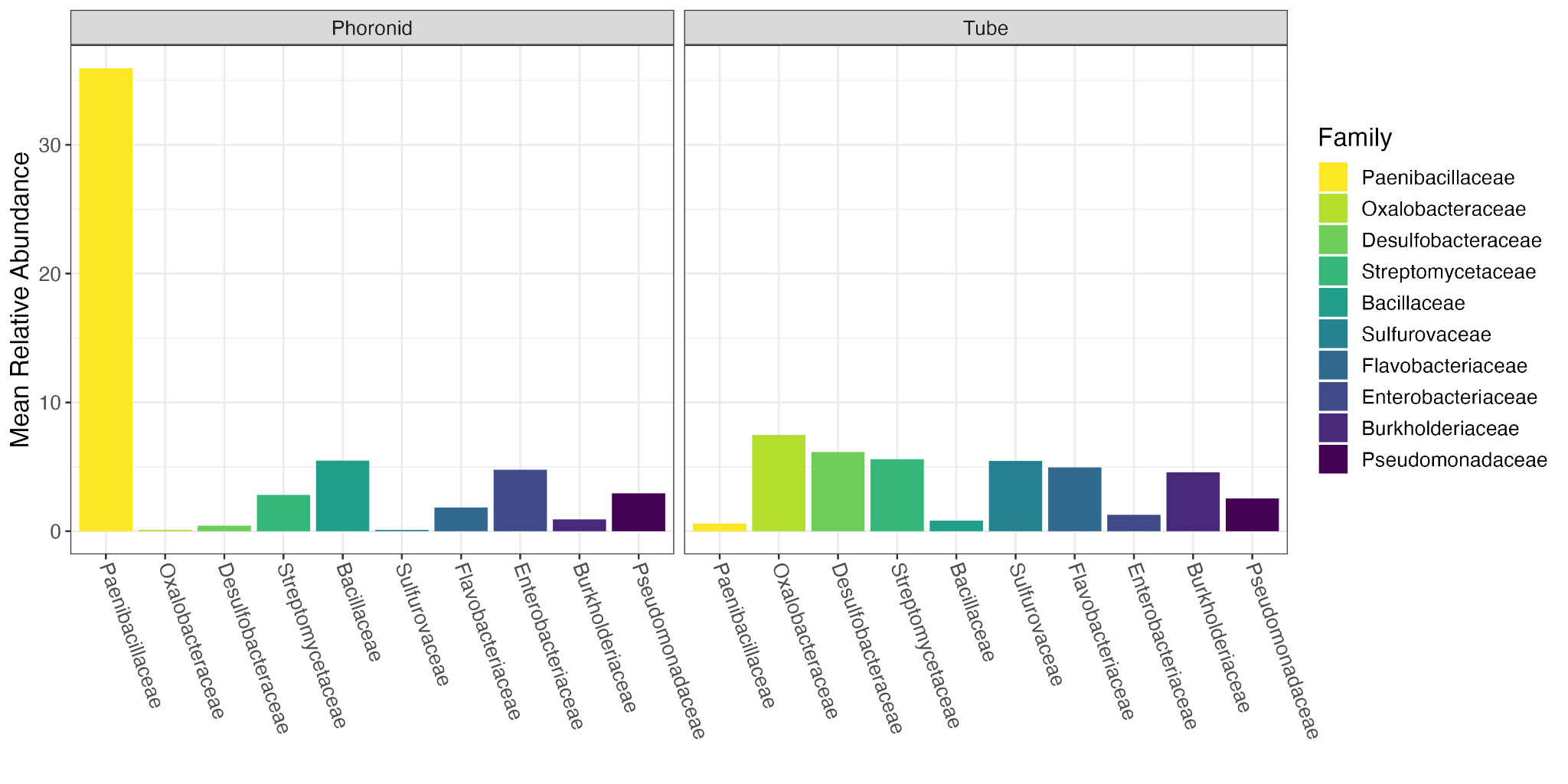

**Figure S3.** Vegetation status plays a role in community structuring. Principal-coordinate analysis (PCoA) visualization of Weighted Unifrac distances shows individual samples designated as points with colors and shapes representing vegetation status: true (gray triangle) or false (black circle). Ellipses represent the 95% confidence interval around the centroid of each group. Number of samples summarized in (A) and (B) as follows - Vegetated: *n_true_*= 25, *n_false_* = 4.

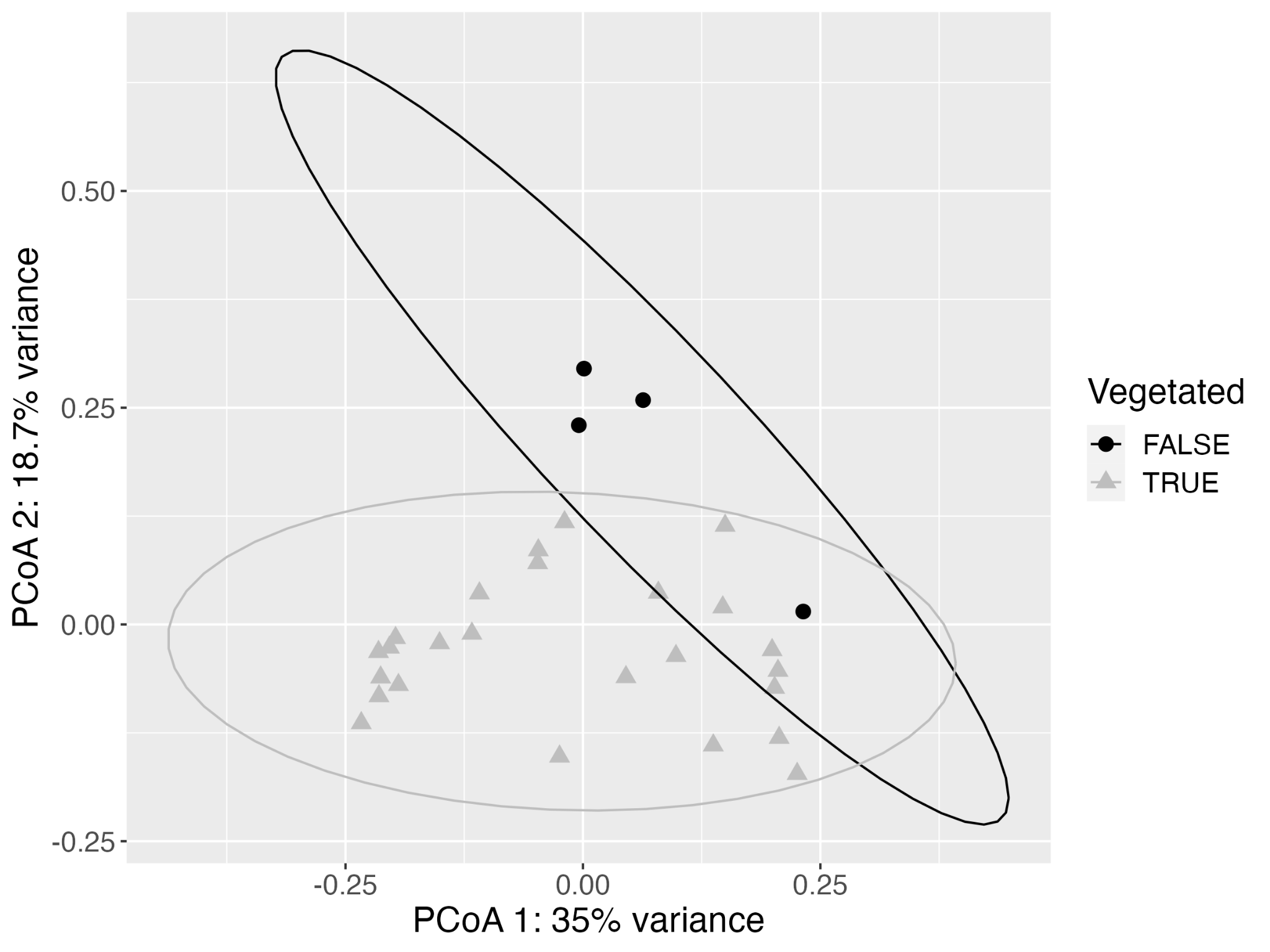

**Figure S4.** METABOLIC gene-level results. Heatmap showing whether the metagenomic coassembly or individual MAGs had any genes detected by METABOLIC related to each of the different sulfur cycling metabolic processes.

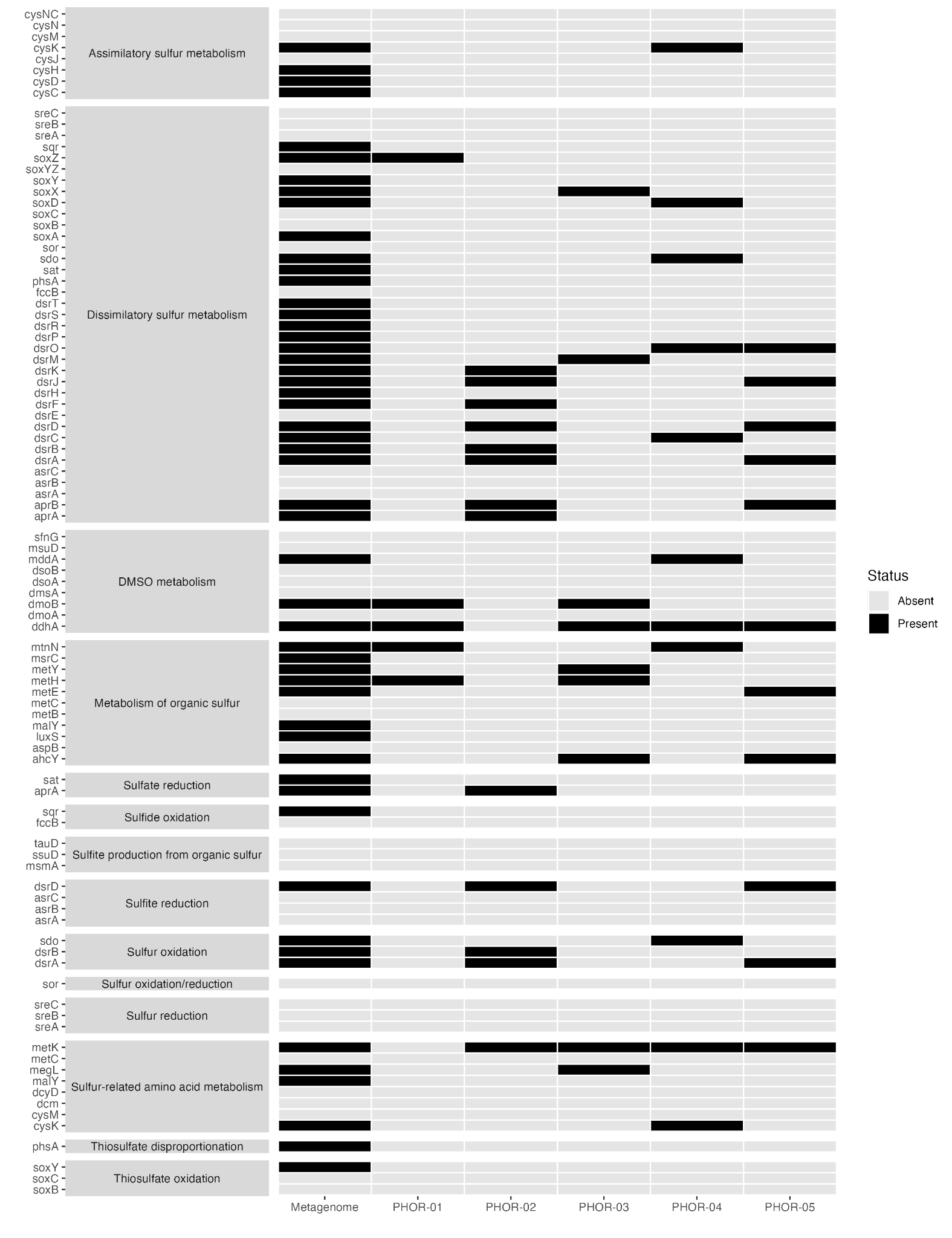

**Table S1.** Taxonomic placement of MAGs by GTDB-Tk. Full taxonomic assignment of each MAG from GTDB-Tk, as well as the calculated relative evolutionary divergence (RED) value of novelty.

| **Bin identifier** | **Phylum** | **Class** | **Order** | **Family** | **Genus** | **Relative Evolutionary Divergence** |
| --- | --- | --- | --- | --- | --- | --- |
| PHOR-01 | Campylobacterota | Campylobacteria | Campylobacterales | Sulfurovaceae | *Sulfurovum* | 0.96 |
| PHOR-02 | Proteobacteria | Gammaproteobacteria | SZUA-229 | SZUA-229 | *GCA-2746365* | 0.95 |
| PHOR-03 | Bacteroidota | Bacteroidia | Flavobacteriales | Flavobacteriaceae | *Aureibaculum* | 0.92 |
| PHOR-04 | Campylobacterota | Campylobacteria |  |  |  | 0.45 |
| PHOR-05 | Desulfobacterota | Desulfobacteria | Desulfobacterales | Desulfosarcinaceae |  | 0.82 |
